# Low-intensity sub-threshold rhythmic 10Hz median nerve stimulation modulates mu-band EEG oscillations in individuals with Tourette syndrome

**DOI:** 10.64898/2026.02.07.704541

**Authors:** Barbara Morera, Ciara McCready, Stephen R. Jackson

## Abstract

Tourette syndrome (TS) is a neurological disorder characterised by the occurrence of vocal and motor tics. Rhythmic median nerve stimulation (MNS) at 10Hz has been shown to cause a substantial reduction in tic frequency in individuals with Tourette Syndrome. The mechanism of action is currently unknown but has been hypothesised to involve entrainment of cortical oscillations within the sensorimotor cortex linked to the initiation of movement. An important methodological detail of these studies is that MNS is delivered at or above threshold (i.e., the minimum stimulation level required to elicit a visible muscle twitch). This is important issue as it means that the observed effects of rMNS could be driven primarily by afferent signals in response to stimulation, the re-afferent signals arising from the muscle, or a combination of these signals. To examine this further, we used electroencephalography (EEG) to investigate the effect of delivering 1s trains of sub-threshold rhythmic 10Hz MNS in a group of 15 adults with TS compared to a matched group of 20 neurotypical control participants. The results demonstrate that the EEG response (somatosensory evoked potential (SEP) to rMNS increased linearly with increasing stimulation amplitude. This was paralleled by substantially increased inter-trial coherence (ITC) during rMNS. Importantly, the duration of increased ITC was reduced for the TS group compared to controls. Importantly, these results were largely similar when analyses were restricted only to sub-threshold trials in which no visible muscle twitch was elicited by MNS. These results confirm that sub-threshold rhythmic MNS is sufficient to modulate somatosensory physiology and may also be sufficient to elicit the clinical benefits previously observed for MNS.

## Introduction

Tourette syndrome (TS) is a neurological disorder of childhood onset that is characterised by the occurrence of vocal and motor tics (Cohen et al., 2013). Tics are involuntary, rapid, abrupt, repetitive, recurrent, and non-rhythmic movements or vocalizations. TS or chronic tic disorder (CTD) affects ∼1-2% of children and adolescents and is the most common form of movement disorder in children (Cohen et al., 2013). The pathophysiology of TS has been linked to dysfunction within cortical-striatal-thalamic-cortical (CSTC) brain circuits involved in the selection and initiation of movements (Albin & Mink, 2006). Specifically, TS has been associated with a reduction in the number of inhibitory fast-spiking parvalbumin interneurons in the striatum (Kalanithi et al., 2005). and it has been proposed that this might lead to increased neural ‘noise’ within the striatum, and in downstream thalamo-cortical sensorimotor networks (Albin & Mink, 2006; Gialopsou et al., in 2025).

Rhythmic median nerve stimulation (MNS) has been shown to result in frequency specific increases in the amplitude and phase synchronisation of neural oscillations (Houlgreave et al., 2022; Morera Maiquez et al., 2020). This modulation is restricted to the contralateral sensorimotor cortex and is not seen during arrhythmic stimulation with an identical number of stimuli and the same mean frequency (Houlgreave et al., 2022; Morera Maiquez et al., 2020). Furthermore, key neurometabolite levels in the sensorimotor cortex (i.e., GABA and Glutamate) have been shown to be modulated by both rhythmic and arrthythmic 10Hz trains of MNS (Houlgreave et al., 2025). Specifically, during periods of 10Hz MNS GABA concentrations are significantly reduced (Houlgreave et al., 2025).

The mechanisms underpinning rhythmic MNS are of therapeutic interest as, compared to periods of no stimulation, rhythmic 10Hz MNS has been shown to produce a substantial reduction in tic frequency and the urge-to-tic in individuals with Tourette Syndrome (Iverson, Arbuckle, Song, et al., 2023; Iverson, Arbuckle, Ueda, et al., 2023; Maiquez et al., 2023; Morera Maiquez et al., 2020). For this reason, and in contrast to other non-invasive stimulation methods such as transcranial magnetic stimulation (TMS), rhythmic MNS offers an attractive approach to modulating brain sensorimotor networks that might easily be developed as a wearable therapeutic device suitable for use outside of the clinic (Maiquez et al., 2023).

An important mechanistic question remains to be investigated, however, as in the above studies, rhythmic MNS was delivered at or above the minimum threshold required to induce a visible muscle twitch. One important consequence of delivering MNS at or above threshold is that, in addition to producing an afferent signal to the somatosensory cortex, the stimulation will also induce a re-afferent signal from the muscle (i.e., there will be a signal to the somatosensory cortex arising from the muscle twitch induced by MNS). It remains to be determined therefore if sub-threshold MNS, that does not induce a visible muscle twitch, remains effective in modulating the cortical brain oscillations linked to movement initiation. In addition to being of theoretical importance, this question is also of considerable clinical importance as sub-threshold stimulation is considered to be far more comfortable that stimulation at or above threshold, particularly in the case of young children who may be put off receiving effective therapeutic MNS if it is delivered at or above threshold.

## Methods

### Participants

Data were collected from 15 participants (10 males) with a confirmed diagnosis of Tourette syndrome or chronic tic disorder, with an age range of 18-53, and an average age of 33.8 (± 12.88) years (see Table 1). A control group consisting of 20 neurotypical adults (8 males) were also recruited, with an age range of 19-59 and an average age of 28.25 (± 10.01) years. Statistical comparison using an independent-samples t-test confirmed that the mean ages of the two groups did not differ (t(33) = 1.45, p = 0.16).

**Table 1.**
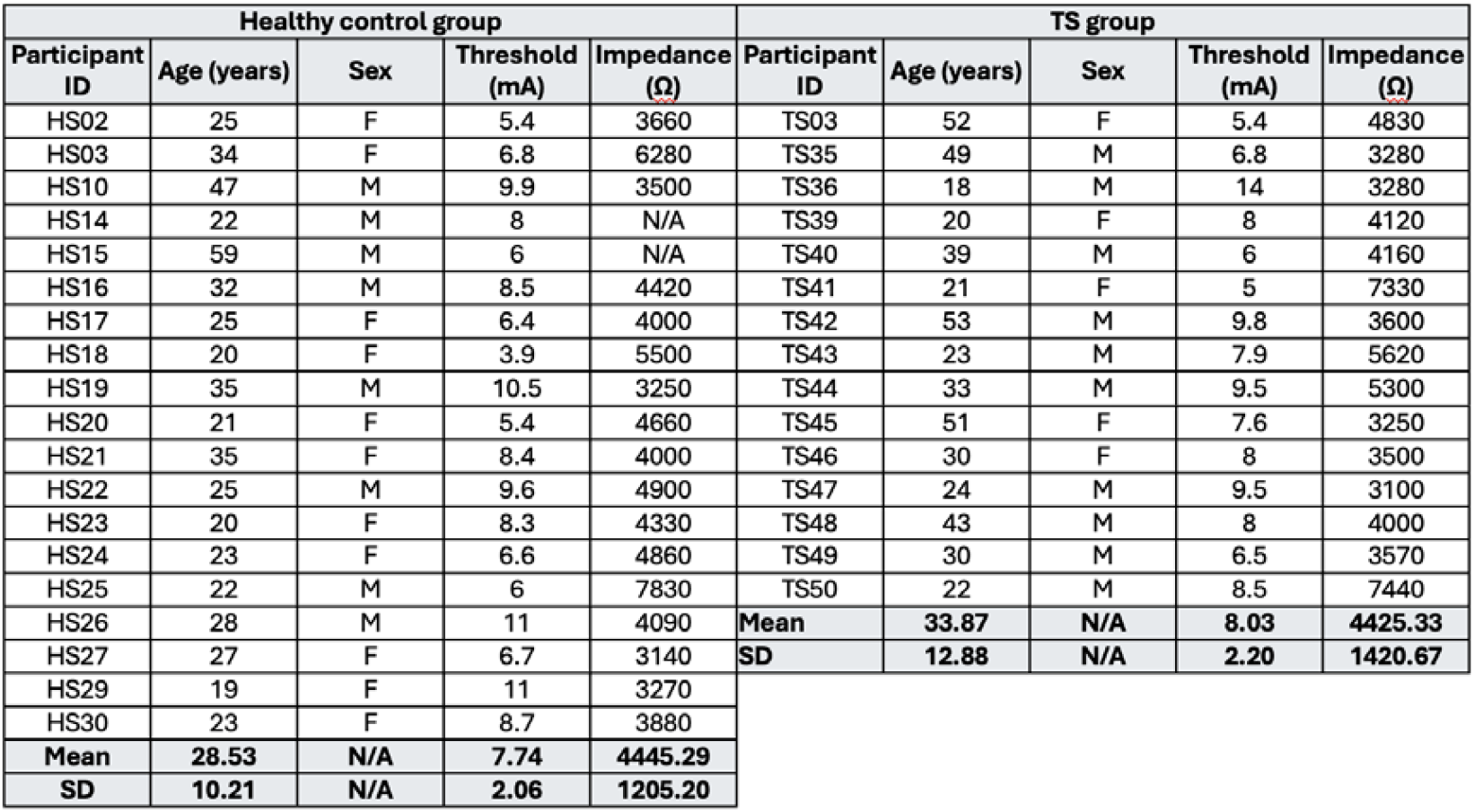
Characteristics of TS and control groups.

| Healthy control group |  |  |  |  | TS group |  |  |  |  |
| --- | --- | --- | --- | --- | --- | --- | --- | --- | --- |
| Participant ID | Age (years) | Sex | Threshold (mA) | Impedance ($\Omega$) | Participant ID | Age (years) | Sex | Threshold (mA) | Impedance ($\Omega$) |
| HS02 | 25 | F | 5.4 | 3660 | TS03 | 52 | F | 5.4 | 4830 |
| HS03 | 34 | F | 6.8 | 6280 | TS35 | 49 | M | 6.8 | 3280 |
| HS10 | 47 | M | 9.9 | 3500 | TS36 | 18 | M | 14 | 3280 |
| HS14 | 22 | M | 8 | N/A | TS39 | 20 | F | 8 | 4120 |
| HS15 | 59 | M | 6 | N/A | TS40 | 39 | M | 6 | 4160 |
| HS16 | 32 | M | 8.5 | 4420 | TS41 | 21 | F | 5 | 7330 |
| HS17 | 25 | F | 6.4 | 4000 | TS42 | 53 | M | 9.8 | 3600 |
| HS18 | 20 | F | 3.9 | 5500 | TS43 | 23 | M | 7.9 | 5620 |
| HS19 | 35 | M | 10.5 | 3250 | TS44 | 33 | M | 9.5 | 5300 |
| HS20 | 21 | F | 5.4 | 4660 | TS45 | 51 | F | 7.6 | 3250 |
| HS21 | 35 | F | 8.4 | 4000 | TS46 | 30 | F | 8 | 3500 |
| HS22 | 25 | M | 9.6 | 4900 | TS47 | 24 | M | 9.5 | 3100 |
| HS23 | 20 | F | 8.3 | 4330 | TS48 | 43 | M | 8 | 4000 |
| HS24 | 23 | F | 6.6 | 4860 | TS49 | 30 | M | 6.5 | 3570 |
| HS25 | 22 | M | 6 | 7830 | TS50 | 22 | M | 8.5 | 7440 |
| HS26 | 28 | M | 11 | 4090 | Mean | 33.87 | N/A | 8.03 | 4425.33 |
| HS27 | 27 | F | 6.7 | 3140 | SD | 12.88 | N/A | 2.20 | 1420.67 |
| HS29 | 19 | F | 11 | 3270 |  |  |  |  |  |
| HS30 | 23 | F | 8.7 | 3880 |  |  |  |  |  |
| Mean | 28.53 | N/A | 7.74 | 4445.29 |  |  |  |  |  |
| SD | 10.21 | N/A | 2.06 | 1205.20 |  |  |  |  |  |

### Median nerve stimulation

MNS was delivered over the dominant (right) wrist, with the electrodes placed over the median nerve (cathode proximal) using a Digitimer DS7A (Digitimer Ltd, UK) device. The stimulation pulse width was set at 0.2ms and the maximum voltage (Vmax) set to 400 V. The motor threshold (MT) was measured individually for each participant and was defined as the minimum MNS intensity required to produce a visible thumb twitch (mean MT (± SD): HC group = 7.68 (2.02) mA; TS group = 8.03 (2.2) mA). Statistical comparison using an independent-samples t-test confirmed that the mean threshold values of the two groups did not differ (t(33) = 0.5, p = 0.62).

### 10 Hz MNS protocol

During each trial MNS was delivered as 10Hz trains of 10 biphasic pulses for a period of one second. Each pulse had a 0.2ms pulse width, with a biphasic ratio of 30%. The gap between the cathodic and anodic pulses was 0.1ms. The stimulation intensities delivered in this study were: 3.5 mA, 4 mA, 4.5 mA, 5 mA, 5.5 mA, 6 mA, 6.5 mA, and 7 mA, all of which fall within the low-intensity range observed in a previous clinical study (Maiquez et. al., 2023).

Each trial consisted of a one second train of 10 pulses followed by a period of 5 seconds of no stimulation. 50 trials were delivered at each of the above intensities (i.e., a total of 400 trials), with the trials presented in a random order. Participants were given a break after every 100 trials (i.e., 10min).

### Electroencephalograpy (EEG) data collection

EEG measurements were obtained utilizing a 64-channel actiCHamp Plus system that incorporated active electrodes. EEG was recorded using Brain Vision Recorder software (Brain Products, UK) with a sampling frequency of 1000 Hz. The impedance of the electrodes was maintained below 30 μV for all the participants. The electrodes placed over the left and right mastoid (TP9 and TP10) were used as reference. Data were filtered using a 45 Hz low-pass and a 1 Hz high-pass filter to remove high-frequency and low-frequency noise, respectively. Noisy channels were identified visually and either deleted or interpolated. Interpolation used data from surrounding channels to estimate the activity at the bad channel, thereby maintaining the dimensionality of the EEG data. After interpolating noisy channels, the data were re-referenced to the linked mastoids and down sampled to 128 Hz.

#### EEG data analysis

EEG data analysis focused on electrode CP3, positioned over the left sensorimotor cortex. EEG recorded at the scalp may contain a mixture of signals arising from different sources, including electrical signals arising from muscle, or resulting from eye-blinks. This is a particularly important consideration to keep in mind when recording EEG from individuals with tic disorders like Tourette syndrome who may exhibit eye and facial motor tics. All EEG signals were inspected for evidence of eye-blinks or muscle artefact (see below). In addition, we conducted an Independent Component Analysis (ICA) of the EEG data to identify and remove muscle and ocular artefacts. ICA is a method of blind source separation that separates a mixture of signals into a number of statistically independent components. A common use for ICA is to identify and remove components associated with ocular and muscle artefacts. We used the Multiple Artifact Rejection Algorithm (MARA) to automatically reject artefacts by classifying them based on six features from spatial, spectral, and temporal domains.

To examine brain activity associated with MNS, epochs of four second duration were extracted that were time-locked (i.e., -1 to 3 seconds) to the first pulse of the MNS pulse train. Epochs with excessive noise were rejected based on the following criteria: signal amplitude exceeding ±100 μV in one or more channels; signal slopes exceeding 50 μV in one or more channels; amplitudes exceeding five times the standard deviation of the probability distribution; a kurtosis statistic larger than five times the standard deviation of the data. Processed data were standardized using z-transformation.

Inter-trial coherence (ITC) was computed to quantify the consistency of 10Hz oscillations at the same phase and rhythm across trials, serving as a measure of phase alignment or phase synchrony. 10 Hz inter-trial phase coherence was calculated using the Matlab *phasecoher* function which computes inter-trial phase coherence for a dataset using Gaussian wavelets.

## Results

### Preliminary comparison of group characteristics

Initial analyses confirmed that the mean ages of the TS group and control group did not differ significantly (TS group mean = 33.9 ± 12.9 years, HC group = 28.5 ± 10.2; t(32) = 1.35, p = 0.19) and the mean simulation thresholds did not differ (TS group mean = 8.03 ± 2.2 years, HC group = 7.74 ± 2.1; t(32) = 0.4, p = 0.69).

### Analysis of inter-trial 10 Hz coherence (ITC) of somatosensory evoked potentials

Mean ITC values were computed for each individual for the time period spanning - 1000ms prior to the onset of the MNS pulse train to 3000ms after MNS onset. Mean ITC values for the period -500 ms to + 2000 ms are presented in Figure 1A for each group. For clarity, only data for stimulation intensity values 4 mA, 5 mA, 6 mA and 7 mA are presented in Figure 1A. Inspection of this figure indicates that ITC values increase substantially in response to the onset of MNS and that the magnitude of this increase is proportional to stimulation intensity level. Furthermore, the magnitude of the ITC response to MNS clearly declines over the duration of the stimulation. Importantly, the data for the TS group and HC group look highly similar.

**Figure 1.**
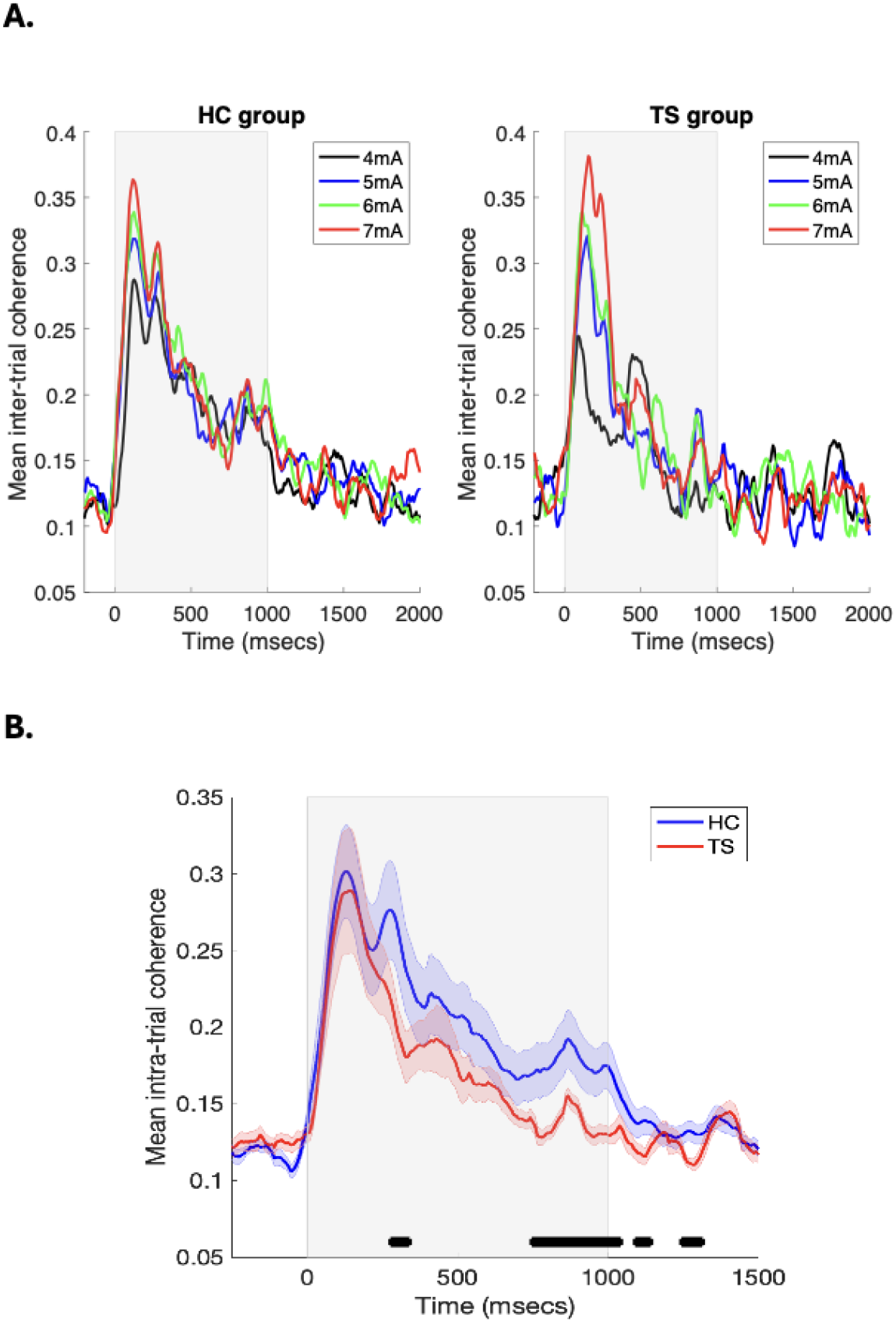
**A**. Shows mean 10Hz inter-trial coherence (ITC) values for the HC and TS groups in response to a one second duration train of 10Hz rhythmic MNS delivered at different intensities (4-7mA). The shaded grey area indicates the period during which stimulation was delivered. **B.** Shows mean ITC values collapsed across stimulation intensity values for each group. The shaded grey area indicates the period during which stimulation was delivered and black symbols indicate periods in which the between-group difference was substantive (effect size >= 0.5).

In order to investigate the effect of stimulation intensity level on the magnitude of the ITC response to MNS, we calculated for each participant and for each level of stimulation intensity (i.e. 3.5 mA to 7 mA) the peak coherence value in the initial 500 ms period after MNS onset. These data were then entered into a linear regression/correlation analysis in which stimulation intensity was used to predicted EMG 10 Hz coherence values. Separate linear regression analyses were conducted for the TS and HC groups. The results of these analyses are presented in Table 2.

**Table 2.** Degree of association (Pearson correlation coefficient) for each group between stimulation intensity level and magnitude of increase in 10 Hz coherence.

|  | **r-value** | **p-value** | **Slope** | **R-squared** |
| --- | --- | --- | --- | --- |
| **HC group** | 0.9 | 0.002 | 0.067 | 0.81 |
| **TS group** | 0.77 | 0.03 | 0.024 | 0.59 |

Statistical comparison between the z-transformed r values presented in Table 2 indicated that these values did not differ between the two groups (p = 0.24).

Mean ITC values collapsed across stimulation intensity values are presented for each group in Figure 1B. Inspection of this figure indicates that ITC initially increases in response to MNS to a comparable level for the TS and HC groups, but the increase in ITC is sustained at a higher level for longer in the HC group compared to the TS group (Effect size: Hedges G >= 0.5).

#### Analysis of the effects of MNS on sub-threshold trials only

In order to examine the effects of 10 Hz pulse trains of MNS on trials in which stimulation was delivered below the minimum level required to elicit a visible muscle twitch (i.e., sub-threshold trials only), we analysed trials for each participant in which stimulation was delivered between 70% and 99% of threshold. Data were collapsed across stimulus intensities for this analysis.

### Analysis of somatosensory evoked potential (SEP) data for CP3

Mean standardised SEP data recorded from the CP3 electrode are presented for each group in Figure 2A. Inspection of this figure confirms that SEP amplitude increases for both groups in response to the onset of MNS, but that this increase is not sustained for the entire duration of the MNS. To investigate this effect further, the data for each individual were segmented into 500 ms epochs (illustrated by the vertical dotted lines in Figure 2A) and the median SEP amplitude was then calculated for each epoch and for each individual. Relevant individual and mean data for each epoch and for each group are presented in Figure 2B.

**Figure 2.**
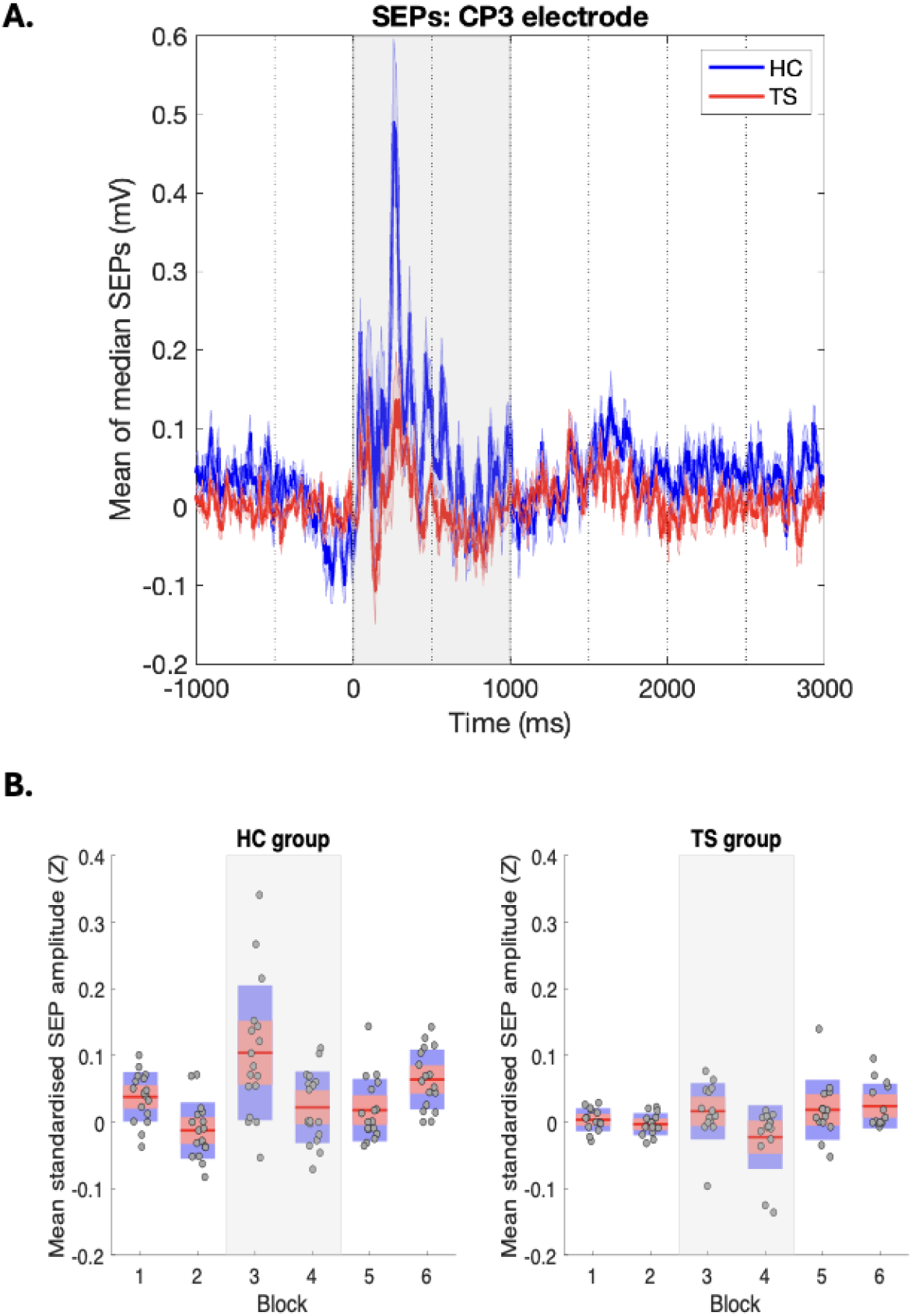
**A**. Shows mean of median evoked responses for the HC and TS groups in response to a one second duration train of sub-threshold 10Hz rhythmic MNS. The shaded grey area indicates the period during which stimulation was delivered. **B.** Boxplots showing median SEPs each 500ms epoch with individual data points shown. The shaded grey area indicates the period during which 10Hz rMNS was delivered.

Statistical comparison of mean data was conducted using a mixed ANOVA with a between subject factor of Group and a within subject factor of Epoch. The ANOVA revealed a significant main effect of Group (F[1,29] = 14.27, p = 0.0007) and Epoch (F[7,203] = 7.95, p = 0.00001) and a significant Group x Epoch interaction (F[7,203] = 3.74, p = 0.0008). Post hoc comparison of group means was conducted at each epoch with false discovery rate correction for multiple comparisons. These analyses revealed that the HC group exhibited increased SEP amplitudes prior to MNS (Epoch 1), during MNS (Epochs 3 & 4), and post stimulation (Epochs 6-8) (minimum t(29) = 2.42, p = 0.03).

#### Time-frequency analysis of EEG data

To further examine this a time-frequency analysis of EEG data using a Continuous Wavelet Transform (CWT) was conducted separately for each group. The results of these analyses are presented in Figure 3. Inspection of this figure indicates that for both groups there is an increase in 5 Hz and 10 Hz EEG power that coincides with the onset of MNS.

**Figure 3.**
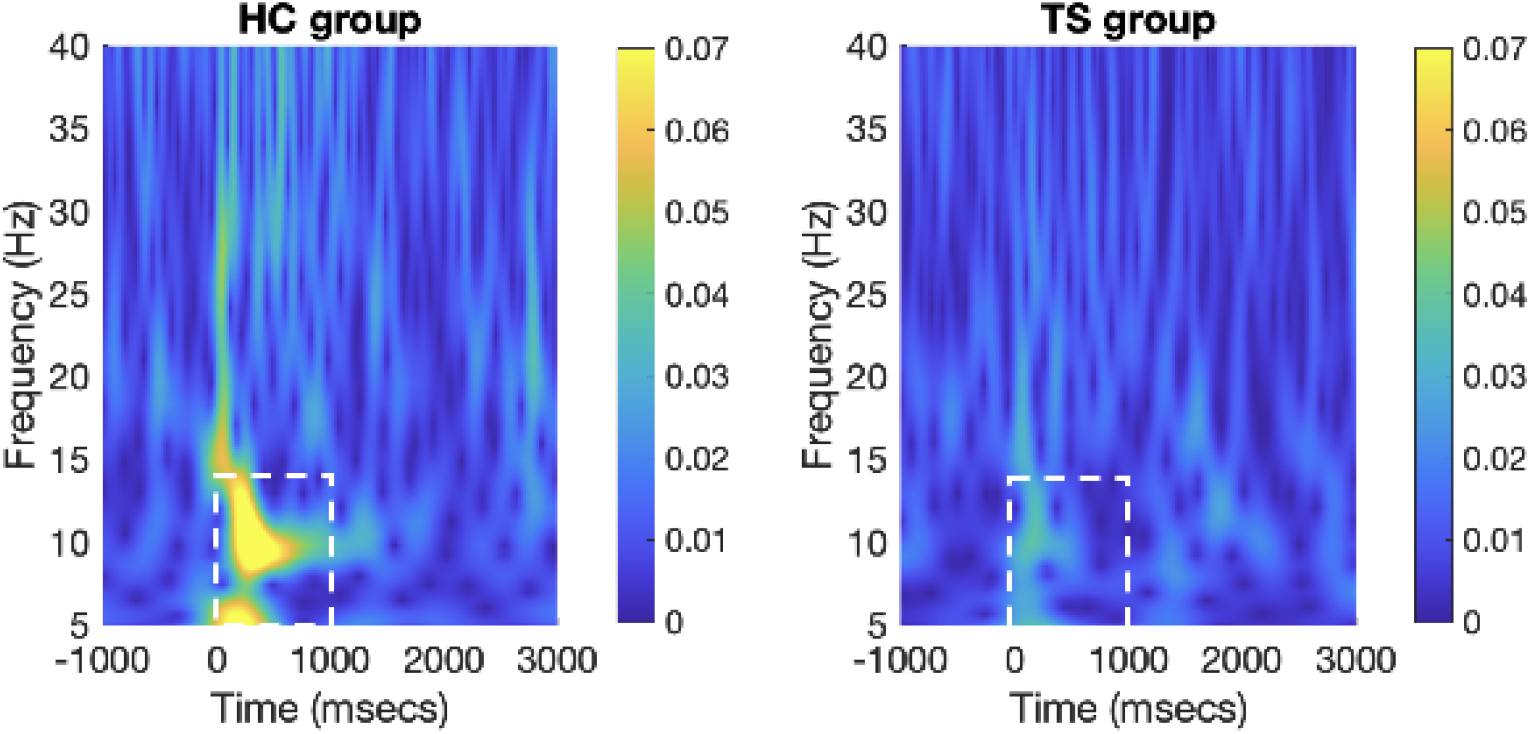
Results of time-frequency analysis of EEG data using a Continuous Wavelet Transform (CWT) for each group. The broken white box indicates the period when 10 Hz rMNS was delivered and the frequency range where modulation of EEG power primarily occurred.

### Analysis of somatosensory evoked potential variability

The variability of SEPs recorded at CP3 was estimated by computing the coefficient of variation (CV) for each epoch and for each individual. The CV is the ratio of the standard deviation and mean (SD/mean). Differences in mean CV were estimated using a mixed ANOVA with a between subject factor of Group and a within subject factor of Epoch. The ANOVA revealed no significant main effect of Group (F[1,29] = 0.9, p = 0.35) and no Group x Epoch interaction (F[7,203] = 1.2, p = 0.3). There was however a significant main effect of Epoch (F[7,203] = 3.59, p = 0.001). This effect was due to an increase in CV at Epoch 3 to coincide the onset of MNS.

### Analysis of 10 Hz inter-trial coherence (ITC) of somatosensory evoked potentials

To examine alterations in inter-trial 10 Hz coherence due to MNS for SEPs recorded at CP3 we computed mean ITC values for each epoch and for each individual. Group means are presented in Figure 3. Differences in mean ITC values were estimated using a mixed ANOVA with a between subject factor of Group and a within subject factor of Epoch. The ANOVA revealed no significant main effect of Group (F[1,29] = 0.9, p = 0.35) and no Group x Epoch interaction (F[7,203] = 1.5, p = 0.16). There was however a significant main effect of Epoch (F[7,203] = 21.1, p < 0.00001). Inspection of Figure 3 indicates that due to an increase in ITC for both groups in response to MNS.

## Discussion

This study used EEG recording to investigate the effects of delivering one second trains of 10 Hz rhythmic median nerve stimulation (MNS) on somatosensory evoked potentials (SEP) in a group of individuals with Tourette syndrome (TS) or chronic tic disorder (CTD) compared to a neurotypical control group. Stimulation was delivered across a range of relatively low stimulation intensities (3.5 – 7 mA) as a key objective for this study was to investigate the effectiveness of low intensity MNS. Key predictions were that 10 Hz rhythmic MNS would increase 10 Hz inter-trial coherence (ITC), that EEG response to MNS would increase with stimulation intensity, and that EEG modulation would be still observed for sub-threshold stimulation.

The main findings from this study can be summarised as follows. 10 Hz ITC values increased substantially in response to rhythmic 10 Hz MNS, and this effect was observed for both the TS group and the controls. Importantly, the magnitude of this increase in ITC was proportional to the level of stimulation intensity delivered. Thus, there was a strong positive correlation between the MNS-induced increase in ITC and stimulation intensity, and the strength of this correlation did not differ between groups. Of interest, the magnitude of the MNS-induced increase in ITC decreased over the course of the period of stimulation and this decrease was greater in the TS group compared to the controls.

An important focus of this study was to determine if rhythmic MNS, delivered at low intensities and specifically, below the minimum threshold required to induce a visible muscle twitch, was nonetheless effective in driving afferent modulation of cortical somatosensory EEG. Analysis of sub-threshold MNS trials only (i.e., 70% - 99% of threshold) revealed that the evoked EEG response to MNS was clearly present in the TS group but was significantly larger in the control group. This effect was confirmed by spectral analysis which confirmed smaller increases in 10Hz power in response to MNS in the TS group compared to controls. This finding of reduced response to MNS in TS relative to neurotypical controls is consistent with the proposal that TS is characterised by increased neural noise in brain sensorimotor networks (Albin & Mink, 2006) and has been operationalised as decreased inter-trial consistency in neural response to stimulation (Gialopsou et al., in 2025). It is also consistent with the finding that resting motor threshold for transcranial magnetic stimulation (TMS) is increased in individuals with TS, particularly children and adolescents (Orth et al., 2008; Pépés et al., 2016). For clarification, this indicates that in general, increased levels of TMS intensity are required for individuals with TS to produce the same magnitude of EMG response to that obtained in neurotypical controls at a lower intensity.

Interest in MNS as a potential non-invasive therapeutic intervention for tic disorders has been recently stimulated by the demonstration that trains of low frequency (alpha-band) MNS can substantially reduce the occurrence of tics, and the urge-to-tic, in individuals with TS (e.g., Morera Maiquez et al., 2020; Iverson, Arbuckle, Song, et al., 2023; Iverson, Arbuckle, Ueda, et al., 2023; Maiquez et al., 2023). Importantly, in each of these studies MNS was delivered using stimulation intensities above the threshold which might contribute to feelings of discomfort or mild and transient skin irritation in a small minority of individuals. The demonstration in the current study that lower intensity, sub-threshold, MNS is sufficient to achieve a comparable neural response to above-threshold stimulation may be clinically important as it may permit rMNS to be effectively delivered at a lower (sub-threshold) intensity ensuring a more comfortable patient experience and reducing the risk of mild transient skin irritation.

## Conclusion

Rhythmic median nerve electrical stimulation (rMNS) delivered at low intensities, specifically at sub-threshold intensities, is sufficient to increase the power and phase-synchrony of cortical EEG responses linked to the initiation of movement in both individuals with Tourette syndrome and a group of matched neurotypical controls. This finding has important clinical implications as it indicates that the therapeutic use of rMNS might proceed using sub-threshold stimulation intensities associated with increased tolerability and reduced levels of discomfort, and reduced likelihood of transient mild skin irritation.

## Acknowledgements

Stephen Jackson is supported by research grants from Medical Research Council (T032588 and UKRI 527), Parkinson’s UK and by an EPSRC IAA grant.

